# *TISCalling:* Leveraging Machine Learning to Identify Translational Initiation Sites in Plants and Viruses

**DOI:** 10.1101/2025.03.06.641763

**Authors:** Ming-Ren Yen, Chia-Yi Cheng, Ting-Ying Wu, Ming-Jung Liu

**Author notes:** **Corresponding author:** Ming-Jung Liu, Biotechnology Center in Southern Taiwan, Academia Sinica, Tainan, Taiwan; Ting-Ying Wu, Institute of Plant and Microbial Biology, Academia Sinica, Taipei, Taiwan.

## Abstract

The recognition of translational initiation sites (TISs) offers complementary insights into identifying genes encoding novel proteins or small peptides. Conventional computational methods primarily identify Ribo-seq-supported TISs and lack the capacity of systematical and global identification of TIS, especially for non-AUG sites in plants. Additionally, these methods are often unsuitable for evaluating the importance of mRNA sequence features for TIS determination. In this study, we present *TISCalling*, a robust framework that combines machine learning (ML) models and statistical analysis to identify and rank novel TISs across eukaryotes. *TISCalling* generalized and ranks important features common to multiple plant and mammalian species while identifying kingdom-specific features such as mRNA secondary structures and G-contents. Furthermore, *TISCalling* achieved high predictive power for identifying novel viral TISs. Importantly, *TISCalling* provides prediction scores for putative TIS along plant transcripts, enabling prioritization of those of interest for further validation. We offer *TISCalling* as a command-line-based package [https://github.com/yenmr/TIScalling], capable of generating prediction models and identifying key sequence features. Additionally, we provide web tools [https://predict.southerngenomics.org/TIScalling] for visualizing pre-computed potential TISs, making it accessible to users without programming experience. The *TISCalling* framework offers a sequence-aware and interpretable approach for decoding genome sequences and exploring functional proteins in plants and viruses.

## INTRODUCTION

The translation of mRNA into protein is at the heart of gene expression. Ribosomes find the correct translation initiation sites (TISs), which determine the protein-coding potential of mRNA and control timely and accurate protein production in response to developmental and environmental cues. Current annotation based on *in silico* prediction was biased to genes that canonically initiate from AUG sites and encode large proteins with known functional domains (Yandell & Ence, 2012; Kearse & Wilusz, 2017; Hsu & Benfey, 2018). Emerging evidence highlights the prevalence of non-canonical translational events, including those from upstream open reading frames (uORFs), translated regions on non-coding RNAs, and features of translation initiation sites from non-AUG codons in plants and plant viruses (von Arnim *et al*., 2014; Hsu & Benfey, 2018; Fang & Liu, 2023; Wu, HL *et al*., 2024b). Thus, a reliable workflow for annotating TISs is essential for decoding genomes and identifying protein-coding genes.

Ribosome sequencing (Ribo-seq), a technique used to globally profile translating ribosome positions, offers *in vivo* evidence for identifying TISs and open reading frames (ORFs) across genomes (Ingolia *et al*., 2009; Ingolia *et al*., 2012). The application of translation inhibitors enhances the resolution of Ribo-seq by selectively enriching specific types of ribosomes: cycloheximide (CHX) stabilizes ribosomes during initiation and elongation whereas lactimomycin (LTM) predominantly stalls ribosomes around initiation sites (Schneider-Poetsch *et al*., 2010). This specificity makes LTM particularly effective for identifying *in vivo* TISs and inferring ORFs with higher resolution (Schneider-Poetsch *et al*., 2010; Lee *et al*., 2012; Li & Liu, 2020). Bioinformatics tools such as RiboTaper and Count, in-frame Percentage and Site (CiPS) utilize ribosome phasing patterns and the magnitudes of ribosome occupancy in CHX-treated Ribo-seq signals to identify AUG TISs and their corresponding ORFs (Calviello *et al*., 2016; Wu *et al*., 2019; Wu, HL *et al*., 2024a). Additionally, TIS hunter (Ribo-TISH) employs LTM- and/or CHX-treated Ribo-seq datasets to detect both AUG and non-AUG initiation sites and their associated ORFs (Zhang, P *et al*., 2017; Wu, TY *et al*., 2024). While these tools provide valuable insights, they rely entirely on Ribo-seq datasets with two focusing specifically on AUG codons for TIS screening. This dependency may limit their ability to comprehensively capture the translational events across genomes. Additionally, Ribo-seq and proteomics/peptidomics resources remain relatively scarce compared to the extensive resources for RNA-seq and protein-DNA interaction sequencing datasets. Therefore, a bioinformatics tool independent of Ribo-seq datasets would serve as a general and comprehensive means of identifying TISs and ORFs across genomes.

Ribo-seq-independent methods for identifying novel AUG and non-AUG TISs rely on either sequence conservation or prediction models. For the former, the comparative-genomic approach has identified dozens of AUG- and non-AUG TISs and their corresponding ORFs across eudicot plant species, primarily located in the 5’ untranslated regions (5’UTRs) of genes (van der Horst *et al*., 2019). However, the reliance on conservation scores and statistical power limits its ability to identify short and non-conserved TIS-initiated ORFs (Couso, 2015). Alternatively, PreTIS, a linear-regression-based prediction model, utilizes mRNA sequence as input to profile potential AUG and non-AUG TISs in 5’UTRs of human and mouse genes (Reuter *et al*., 2016). While promising, PreTIS was developed using previously identified AUG and non-AUG TIS in the 5’UTRs of human and mouse genes, leaving its applicability to plants and its effectiveness at profiling TISs within CDSs uncertain. Together, the available Ribo-seq-dependent and independent approaches are constrained by their individual limitations and unable to deliver a comprehensive and systematic profile of AUG- and nonAUG-initiated translational events across different genic regions—5’ and 3’UTRs and within CDS regions—or across entire plant transcriptome. Developing more integrative and versatile tools remains a critical challenge for fully decoding translational landscapes.

Here, we introduce *TISCalling*, a sequence-aware, ML-based framework for TIS predictor that enables the identification of key mRNA sequences regulating TIS recognition and the systemic profiling of potential TISs along transcripts (**Fig. 1**). Using publicly available datasets of the *in vivo* TISs identified through Ribo-seq (Lee *et al*., 2012; Willems *et al*., 2017; Li & Liu, 2020), we developed predictive models to identify both AUG and non-AUG TISs in plants (e.g. Arabidopsis and tomato) and mammals (e.g. human and mouse) and analyze the mRNA sequence features that significantly influenced predictive performance. The predictive models were applied to identify potential TIS sites with high prediction scores in UTRs of plant stress-related genes, non-coding RNAs, and even viral genomes. To ensure accessibility and flexibility, we provided the *TISCalling* framework as a command-line-based tool on Github [https://github.com/yenmr/TIScalling], enabling users to identify mRNA features and generate prediction results. Additionally, by integrating TIS models into a user-friendly web tool [https://predict.southerngenomics.org/TIScalling], the *TISCalling* facilitates the visualization of TISs along genes. By providing a sequence-aware approach independent of Ribo-seq datasets, *TISCalling* supports the discovery of AUG and nonAUG TISs and small ORFs, contributing to the decoding of plant and viral genomes.

**Figure 1.**
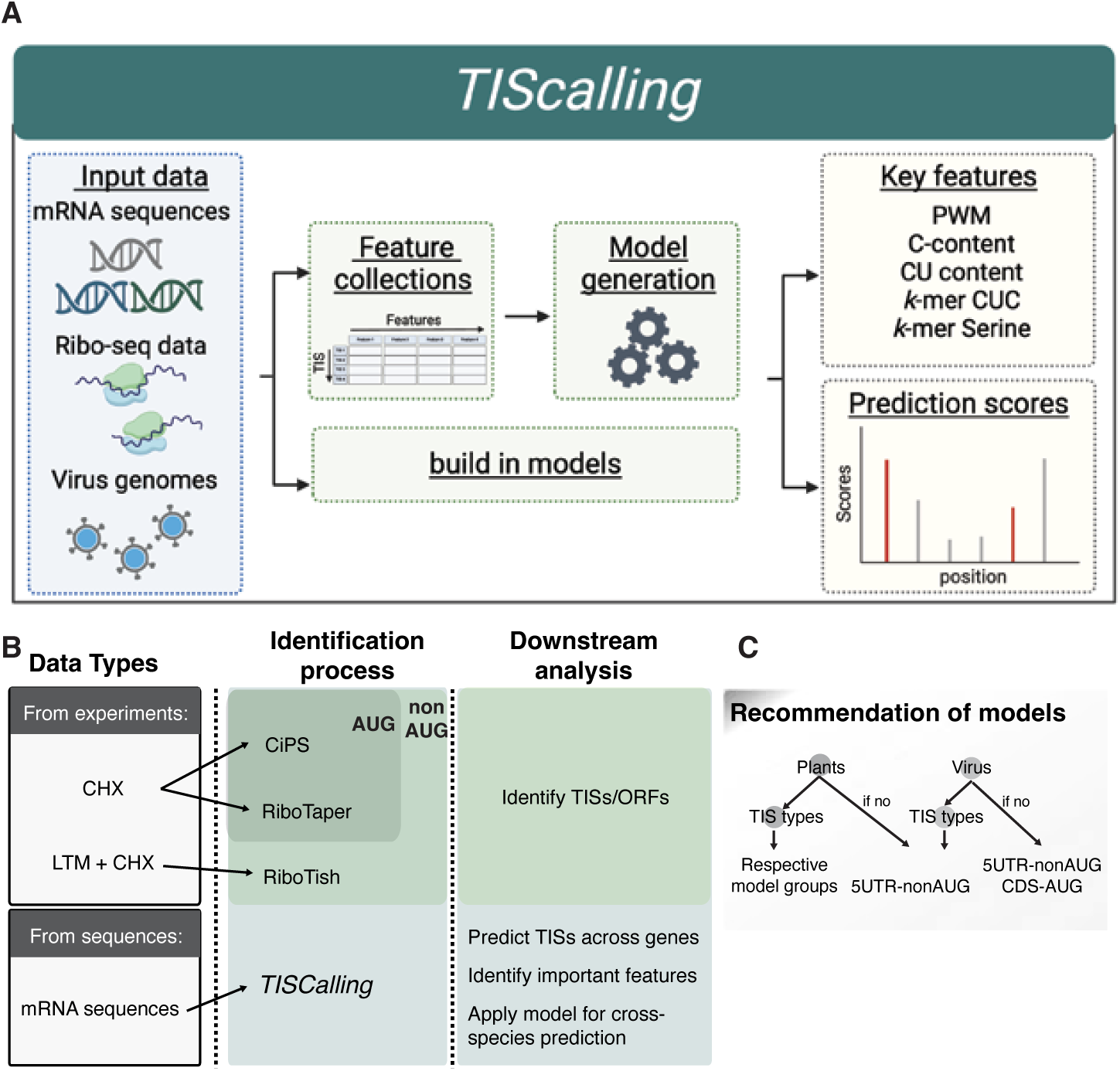
The TISCalling framework, along with its features and applications, is designed for identifying potential TISs in plants and viruses. (A) The TISCalling framework included three parts: the collection of Ribo-seq supported translation initiation sites (TISs) and genome information (blue box), the generation of sequence features and prediction models (green box), and the identification of the key features important for model performance and the application of models for reporting TIS prediction scores (yellow box). Additionally, by inputting the sequences of mRNAs and viral genomes into the build-in models (i.e., the one generated previously), TISCalling can directly report the scores along sequences. (B) The flowchart outlined the required input data types, the pipelines of TIS and open-reading frames (ORFs) identification, and their outcomes. It illustrated the relationship between the types of available input datasets and the corresponding AUG and/or nonAUG TIS/ORF identification methods (indicated by arrows), and the various downstream outcomes that these methods can generate (highlighted in colored boxes). (C) Model recommendations for TIS identification. The flowchart illustrates the recommended predictive models with/without target TIS types for identifying plant and viral TISs.

## MATERIALS and METHODS

### Dataset collection

We collected datasets of the novel (i.e. unannotated) translation initiation sites (TISs) with significant translation initiation activities, referred as to true positive (TP) datasets, from tomato and Arabidopsis LTM-treated ribosome profiling data as reported by Li and Liu (2020). The TP TIS datasets for human HEK293 cells and mouse MEF cells were retrieved from the previous publication (Lee *et al*. (2012). We collected the public TP TIS datasets generated from different plant and virus studies (**Supporting Dataset S1**). Briefly, for Arabidopsis, the novel TIS data included those reported in the study by (Wu, TY *et al*., 2024) and those associated with non-coding open-reading frames (ORFs), downstream ORFs, upstream ORFs (uORFs), and within coding regions (CDSs) as reported by (Wu, HL *et al*., 2024a). In tomato, the novel TISs from uORFs and small ORFs on de novo assembled transcripts were retrieved from Wu *et al*. (2019). For human and plant viruses, novel TIS datasets were sourced from cytomegalovirus (HCMV) (Stern-Ginossar *et al*., 2012), SARS-CoV-2 (Finkel *et al*., 2021), and Tomato yellow leaf curl Thailand virus (TYLCTHV; genus Begomovirus; Chiu *et al*. (2022).

To construct a dataset of true negative (TN) TISs, for each positive TIS in our dataset, we collected both ATG and near-cognate codon sites that were located upstream of the most downstream TP TIS within the same transcript and were not marked as TP TISs as described by (Wu, TY *et al*., 2024). This methodology generated robust TP and TN datasets for assessing the accuracy of our model in distinguishing between actual translation initiation sites and potential false positives.

### Feature Collection and Dataset Balancing

We focused on the mRNA sequences information around the TP and TN TISs and gathered 1,250 features for each TIS, which were segmented into three categories: 5 known features, such as the Kozak sequence, TIS codon usage and adjacent flanking sequences (Kozak, 1984; Kozak, 1989; Noderer *et al*., 2014; Reuter *et al*., 2016; Zhang, S *et al*., 2017; Diaz de Arce *et al*., 2018; Li & Liu, 2020), 18 ORF features that included mononucleotide contents, secondary RNA structures within or upstream of ORFs, and ORF sizes, and 1,227 contextual features detailing the nucleotide/amino acid frequency of k-mers within a 200-nt region centered on a TIS. Further details on these features are described below.

Feature collection mainly followed the classifications and methodologies outlined in Wu, TY *et al*. (2024). This included categorization and slight modifications to the existing frameworks of known, contextual, ORF and TIS codon usage features. Each category was described below in detail:

#### Position-Weight Matrix (PWM)

PWM-related features were derived to represent the relationship between the flanking sequence context and TIS translational efficiency. Features such as “PWM” were based on a PWM matrix generated from a given TP group’s flanking sequences (positions −15 to +10) with *in vivo* translation initiation activity.

#### Noderer translational efficiency

The feature was derived from (Noderer *et al*., 2014), which analyzed the flanking sequence context from positions −6 to +5 around the AUG translational start.

#### Kozak sequence context

The Kozak sequence context was categorized into two levels: strong Kozak (A or G at −3 and G at +4), and no Kozak context.

#### TIS-codon usage

The TIS codon usage was determined as described previously with small modifications (Zhang, S *et al*., 2017). Briefly, for a given codon, the proportion of the target codon sites among all the identified TP TISs was normalized to the proportion of the target codon sites among all codon sites found in the transcript regions of all annotated genes. The corresponding log_2_ ratio was then computed and referred to as the feature value of TIS codon usage bias. Codons with negative values were excluded.

#### ORF features

Features included the A/T/C/G mononucleotide contents in the upstream regions and within the ORF regions of the TP/TN TIS-initiated ORFs and their ORF lengths. Additionally, features included the distance from the TP/TN TISs to the upstream region of the nearest start and stop codons.

#### Minimum Free Energy (MFE) of mRNA secondary structure

MFEs were calculated using the RNAfold program (Lorenz *et al*., 2011) for 80-bp regions centered on TISs, with a sliding window of 20-nt and a step size of 10-nt. Furthermore, the magnitudes of MFE differences were normalized across nearby regions to highlight the structural dynamics around the TISs.

#### Contextual features

We quantified the frequency of all possible k-mers (k = 1 for position-specific k-mers and k = 3 for codon and respective amino acid k-mers) in a 199-nt window around the TIS site. This included in-frame and out-of-frame k-mers, as well as those upstream and/or downstream of the TIS site. Additionally, the frequency of all possible amino acids with lengths of 1 in the 99-nt regions downstream of the TIS site and within a TIS-initiated ORF was analyzed, contributing to a total of 1,227 contextual features.

To prevent missing values in the features set, input sites with fewer than 100 bp of flanking sequences were padded with ‘N’. The feature values were scaled using the MinMaxScaler from the scikit-learn package, rescaling them from 0 to 1. This scaling process ensures that all features contribute equally to the model, preventing features with larger numerical ranges from dominating others.

### Feature Selection for Model Training and Testing

To have balanced data for model training and evaluation, we downscaled the number of true positives (TPs) and true negatives (TNs) equally via random sampling without replacement. For the division of the dataset into training (80%) and testing (20%) segments, we selected the final feature set based on the correlation between features and the significance of their enrichment between true positives (TPs) and true negatives (TNs). This selection process was guided by research suggesting that an excess of features can degrade model performance (Bzdok & Meyer-Lindenberg, 2018). We employed the Wilcoxon rank-sum test with Bonferroni correction to assess the statistical significance of differences between TP and TN sites. Pearson correlation coefficients were then calculated among the contextual and ORF features. We retained the 50 most significant (those with the smallest adjusted p-values) and uncorrelated (*r* < 0.7) contextual features, as well as the uncorrelated (*r* < 0.7) ORF features with adjusted p-values less than 0.05 for the model training step. To facilitate feature evaluation, Cohen’s d was calculated to show the effect size that quantifies the difference between two group means relative to their standard deviations. In this context, it helps assess how well a feature distinguishes TPs and TNs, with higher values indicating better discriminative power.

### Development and Evaluation of Machine Learning (ML) Models

An ML pipeline was described previously (Uygun *et al*., 2019; Wu *et al*., 2021). Briefly, we used scikit-learn version 1.5.0 in Python version 3.10.14 to train and test the models.

We constructed ML models using four different algorithms to evaluate their efficacy in predicting TISs: Logistic Regression (LR), and Support Vector Machine (SVM). To optimize these models, we employed a grid search for each algorithm to identify the optimal settings, coupled with 5-fold cross-validation. The F1 scores value was used to select the best model for each TIS group. The models with the highest F1 values underwent further testing on a separate validation set to confirm their generalizability and robustness. The hyperparameters tested were as follows: LR: C = [0.01, 0.1, 1, 10, 100], intercept_scaling = [0.1, 0.5, 1, 2, 5, 10], penalty = [“l2”]. SVMlinear: kernel = [“linear”], C = [0.01, 0.1, 1, 10, 100].

The robustness and feature significance of our models were assessed by applying this machine learning workflow across ten randomly balanced TP and TN datasets.

### Generation of TIS prediction scores

The input for TIS prediction was the mRNA sequences in FASTA format, which provided the sequence information of potential TIS sites at AUG and non-AUG codons (i.e., the near-cognate ones) as well as the sequence information of their flanking regions. For a given triplet being inquired, 1,250 features were generated to provide information of specific features required for the selected prediction model to generate its TIS prediction scores. The output included the prediction probability as well as detailed site information, such as gene ID, position, and the sequence surrounding each potential site.

### Construction of JBrowse and Visualization

#### Visualization

the JBrowse 2 was employed to show TIS prediction scores along genes across multiple species, for the genomes of *Arabidopsis thaliana*, and *Solanum lycopersicum* (tomato), (Diesh *et al*., 2023) as well as to visualize the genome/encoded protein sequences and the annotations of genes. The publicly available Ribo-seq datasets with LTM treatment from Arabidpsis and tomato as described previously (Willems *et al*., 2017; Li & Liu, 2020; Wu, TY *et al*., 2024) and the Ribo-seq datasets with CHX treatment from heat-treated pollens (Poidevin *et al*., 2021) were provided.

#### Data Preparation

Potential TISs for each gene were computed, and the coordinates of each predicted TIS were mapped to their respective genomes and saved in BigWig format to enable efficient visualization. Aligned Ribo-seq and RNA-seq reads were converted to BedGraph files. To standardize read depth across samples, coverage profiles were normalized to reads per million (RPM). BigWig files were subsequently generated from the BedGraph files using the bedGraphToBigWig tool from the UCSC binary utilities (Kent *et al*., 2010).

#### JBrowse Configuration

A JSON configuration file was generated to facilitate the display of all data tracks in JBrowse. This configuration included sequence files, annotation files, RNA-seq and Ribo-seq abundance tracks, and tracks for TIS prediction scores. To streamline visualization, tracks from the positive and negative strands were merged. The AUG model reports the AUG triplet with scores > 0.5 whereas the non-AUG model reports the triplet either at AUG or near-cognate codons with scores > 0.8.

## RESULTS

### *TISCalling* framework: machine learning models for identifying key mRNA features and predicting potential TISs across species

To develop a machine-learning (ML) model for predicting TISs across eukaryotic species, we utilized publicly available datasets of *in vivo* AUG and non-AUG TISs from Arabidopsis (*At*), tomato (*Sl*), human (*Hs*) and mouse (*Mm*). These datasets, derived from LTM-seq (Lee *et al*., 2012; Willems *et al*., 2017; Li & Liu, 2020), employed ribosome profiling with LTM treatment, which enriched ribosome stalled at translation initiation sites during the first round of translation (Schneider-Poetsch *et al*., 2010) (**Supporting Dataset S1**). The TISs identified across 5’UTRs and CDS regions were classified into four groups: 5’UTR-AUG, 5’UTR-nonAUG, CDS-AUG, and CDS-nonAUG (Wu, TY *et al*., 2024). Sequence information from the 200-bp regions flanking each TISs was extracted to build predictive models using logistic regression (LR) and support vector machine (SVM) algorithms (**Figs. 1, 2A; Supplemental Fig. S1**; see **Methods**). The prediction models for 5’UTR-AUG, 5’UTR-nonAUG, and CDS-AUG TISs in Arabidopsis and tomato achieved high performances, with AUROC > 0.90. However, the CDS-nonAUG models showed reduced performance, with AUROC > 0.57 (**Fig. 2A; Supporting Dataset S2** for additional performance metrics). Consequently, we focused on the high-performing models of 5’UTR-AUG, 5’UTR-nonAUG, CDS-AUG TISs in the subsequent analyses. We observed comparable results for human and mouse models (**Fig. 2A**). When testing cross-species applicability, models trained on Arabidopsis performed best on predicting tomato TISs, followed by human and mouse TISs (**Fig. 2B**, *At*). Similarly, tomato-derived models performed well on other plant species (**Fig. 2B**, *Sl*). Human and mouse models showed high cross-predictive accuracy with AUROC > 0.73 when applied to each other and > 0.68 for Arabidopsis and tomato TIS (overall mean, **Fig. 2B**, *Hs* and *Mm*). These findings indicate that TIS models are broadly applicable across species, with within-kingdom predictions outperforming cross-kingdom predictions (**Supporting Dataset S2** for full results).

**Figure 2.**
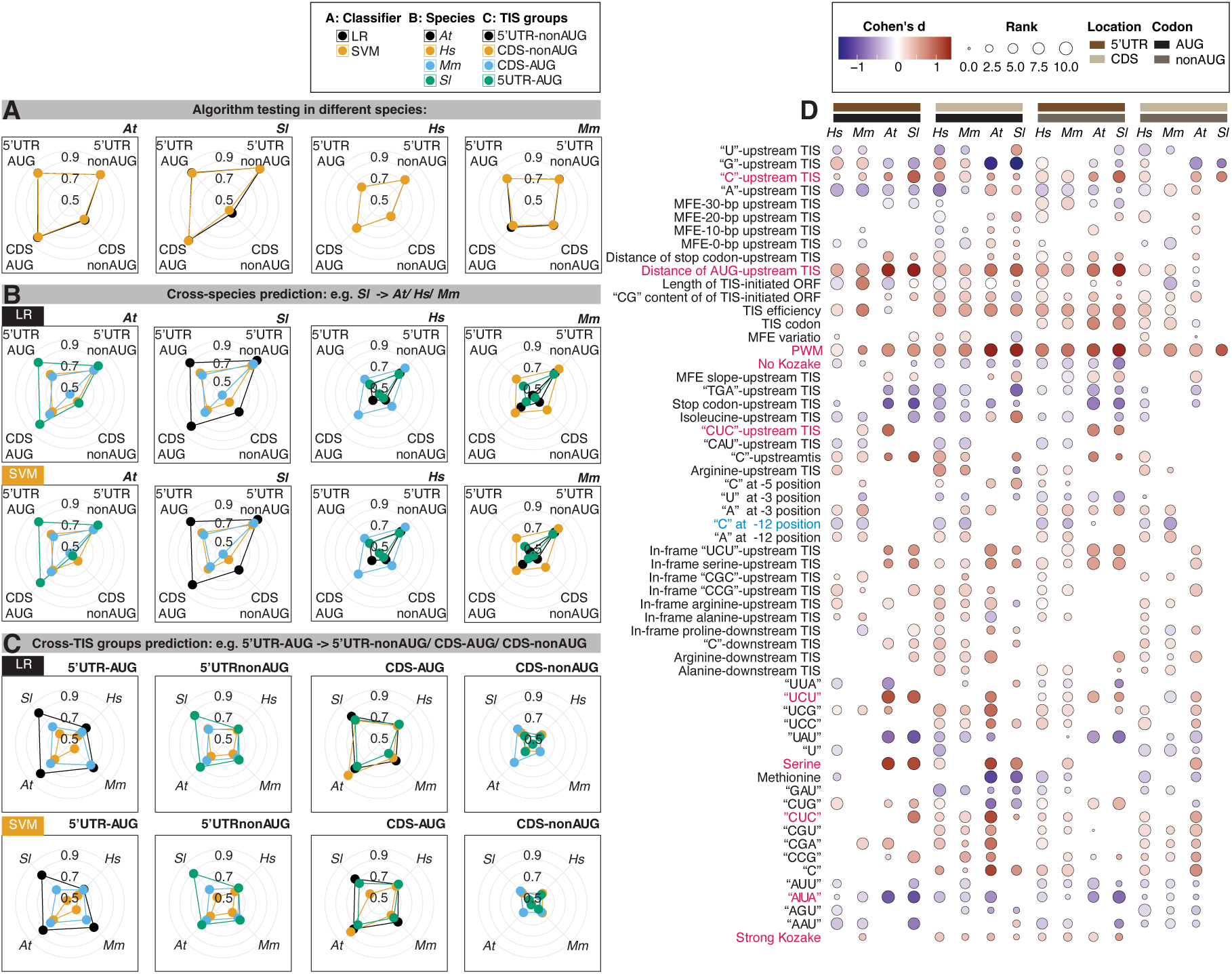
The TISCalling framework-generated prediction models for both AUG and non-AUG TISs across plants and mammals. (A) The performance shown in AUROC scores of the logistic regression (LR) and support-vector-machine (SVM) algorisms in predicting TISs identified from Arabidopsis (At), tomato (Sl), human (Hs) and mouse (Mm). The identified TISs were categorized into four groups based on their locations (i.e., within 5’UTR and main CDS) and the codons of the TIS sites (i.e., AUG and non-AUG codons). (B) Cross-species prediction performance, using models trained on one species to predict TISs in another. The model performance was shown for the four TIS categories as described in panel (A). (C) Cross-TIS prediction performance, using a model trained on one TIS group to predict another TIS group within the same species, as indicated in panel (A). (D) Input features ranked by importance scores derived from the models predicting four types of TISs from different species. Higher ranks indicate greater importance, with features colored according to the difference in mean feature values between true positives (TPs) and true negatives (TNs), as represented by Cohen’s d effect size. Higher effect sizes mean that the mean feature values of TPs are greater than those of TNs.

An analysis of sequence features contributing to model performance revealed several important features shared across TIS models in all four species. These include the preference for PWM features (i.e., the TIS-proximal sequences spanning positions −15 to +10), CU-related sequences, and the avoidance of upstream AUG sites and short AUA sequences near a TIS (features marked in red in **Fig. 2D**, see **Methods** for feature collections and **Supporting Dataset S3** for detailed feature information). Interestingly, the presence of “C” at the −12 position was an important feature for mammalian TIS prediction, consistent with previous findings (Reuter *et al*., 2016), whereas this feature was not significant in plant models, highlighting kingdom-specific features (features marked in blue in **Fig. 2D**).

To investigate the generality of TIS recognition mechanisms across four TIS types, we assessed cross-prediction performance using models derived from one TIS type to predict the remaining three types. Overall, cross-TIS type prediction achieved AUROC values > 0.7 (mean, **Fig. 2C**) but performed less accurately compared to within-TIS type prediction (**Fig. 2A**). For instance, the 5’UTR-nonAUG model from Arabidopsis achieved ~0.94 for predicting 5’UTR-nonAUG TISs in tomato but only ~0.85 when predicting Arabidopsis 5’UTR-AUG TISs. These results suggest a combination of common and specific sequence features associated with different TIS types. For example, the avoidance of upstream AUG sites was observed in both 5’UTR-AUG and 5’UTR-nonAUG TIS groups, Kozak-related features were more prominent in 5’UTR nonAUG TISs (**Fig. 2D**). Together, the *TISCalling* pipeline successfully generated predictive models for AUG and non-AUG TISs in 5’UTRs and CDS regions within and across species. In addition to robust prediction, it revealed conserved and species-specific sequence features underlying TIS recognition, offering novel insights into translational regulation in plants and animals.

### *TISCalling* framework robustly profiles potential TISs in plant transcripts

Our TIS prediction model was developed using TISs experimentally supported from LTM-treated Ribo-seq datasets of Arabidopsis suspension cells and tomato leaves and statistically identified via TIS/ORF-identification methods (**Supporting Dataset S1**) (Lee *et al*., 2012; Willems *et al*., 2017; Machkovech *et al*., 2019; Li & Liu, 2020). To evaluate model robustness across datasets generated using different treatments, we incorporated additional TIS datasets derived from both LTM- and CHX-treated Ribo-seq experiments and diverse identification pipelines (**Supporting Dataset S1**) (Wu *et al*., 2019; Li & Liu, 2020; Hiragori *et al*., 2023). LTM preferentially stalls ribosomes during the first round of translation, providing high-resolution Ribo-seq signals around initiation sites, while cycloheximide (CHX) globally blocks elongating ribosomes across transcripts. Despite these differences, our Arabidopsis 5’UTR-AUG models achieve AUROC > 0.69 when predicting TISs identified using LTM-seq from Arabidopsis seedlings and CHX-seq data from Arabidopsis and tomato seedlings and through RiboTaper and CiPS pipelines (**Fig. 3E**, Known TIS types). Similar results were observed for 5’UTR-nonAUG and CDS-AUG models (**Fig. 3E**, Known TIS types). Strong correlations in prediction performance between Arabidopsis and tomato models (*rho* = 0.95, **Fig. 3E**, Known TIS types) further supported the cross-species applicability of *TISCalling* models in plants (**Fig. 2B**). Moreover, the models successfully identified AUG-TISs within 3’UTRs as well as in non-coding and *de novo*-assembled transcripts, demonstrating their versatility across diverse transcript types (**Fig. 3E**, Others; see **Supporting Dataset S2** for full results).

**Figure 3.**
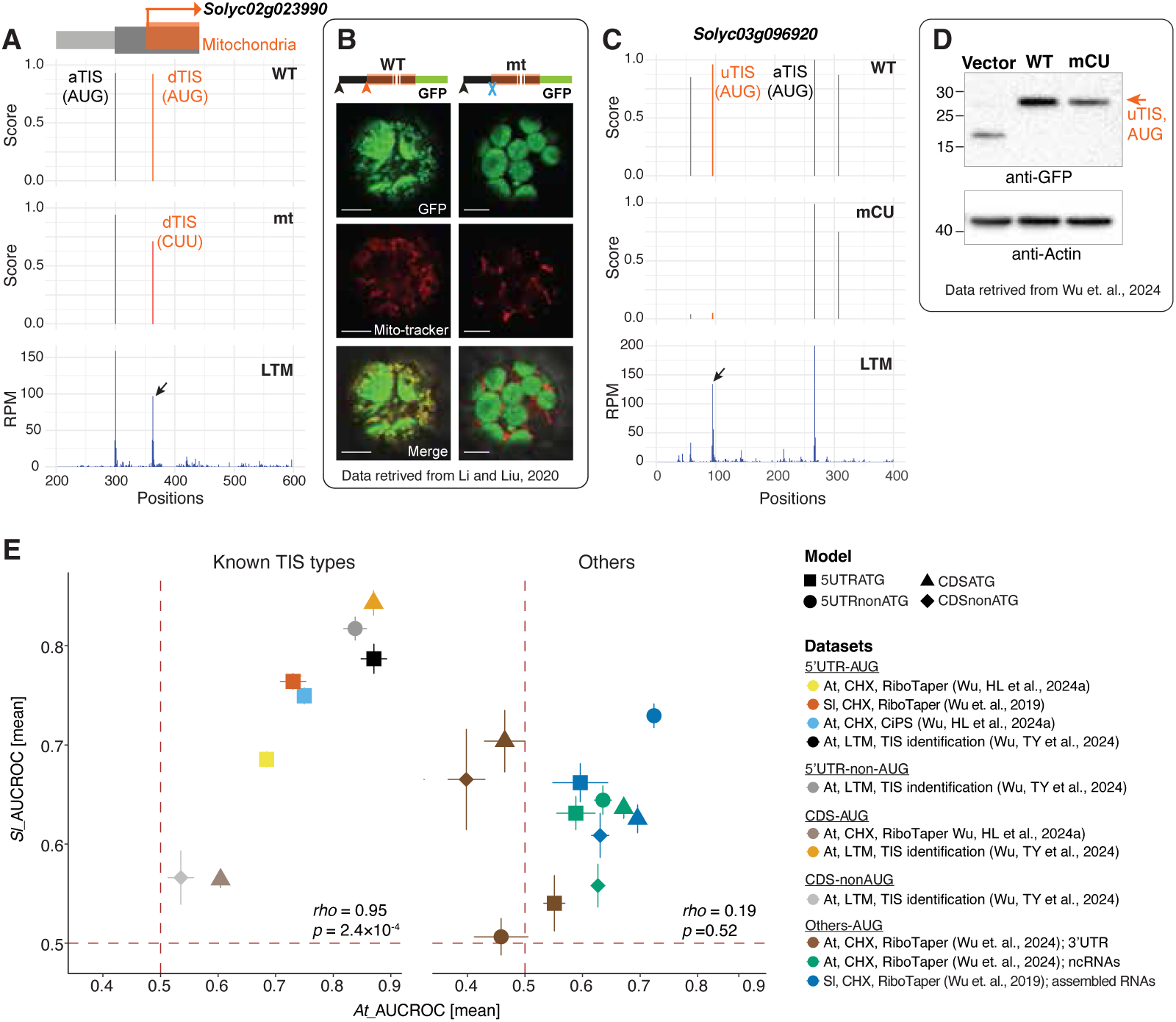
TIS prediction models using flanking mRNA sequences robustly identify plant TISs. (A) The prediction scores of a given triplet along the Solyc02g023990 transcript were generated via the plant CDS-AUG-TIS model and were based on the sequences of the triplet and its flanking 200-bp regions. The AUG triple sites with a prediction score >0.8 along the wild-type (WT) mRNA sequences were shown, including the annotated AUG TIS (aTIS) and a downstream in-frame AUG TIS (dTIS) encoding a mitochondria-localized protein. mt: the AUG dTIS was mutated to CUU triple in the mutant. The LTM plots showed the read density (reads per million mapped reads; RPM) derived from tomato LTM-treated Ribo-seq datasets. In the gene models (top), light and dark gray boxes indicate UTRs and annotated CDSs, while orange boxes indicate dTIS-initiated ORF, which encodes a protein isoform predicted to be localized to mitochondria. (B) Localization of Solyc02g023990-GFP protein with translation driven by the wild-type CDS (left panel) and dTIS-mutated CDS (middle panels). The aTIS, dTISs, and mutated TISs are indicated by a black arrow, orange arrow, and blue cross, respectively. Mitochondria marker: CD3-992 (Nelson et al., 2007). The results were derived from a previous study (Li & Liu, 2020). (C) As in (A), but for the Solyc03g096920 transcript with an upstream AUG TIS (uTIS). mCU: the mutations at CU-rich sequences of the upstream 100-nt region of the uTIS. The results were derived from a previous study (Wu, TY et al., 2024). (D) Expression of the Solyc03g096920-GFP proteins initiated from the uTIS indicated in (C), with translation driven by the upstream 100-nt WT or mCU sequences. Vector: tobacco leaves infiltrated with agrobacteria containing the expression vector (i.e., the GFP plasmid without a target gene sequence). (E) Prediction performance of the four TIS-type models (from tomato (Sl) and Arabidopsis (At), as indicated in Fig. 2A) in predicting TISs derived from previous studies and identified via different TIS/ORF identification pipelines including RiboTaper and CiPS and a TIS identification method (Machkovech et al., 2019; Wu et al., 2019; Wu, HL et al., 2024a; Wu, TY et al., 2024). These previously identified TISs were categorized into two groups: known TIS types including 5’UTR-AUG, 5’UTR-nonAUG, CDS-AUG and CDS-nonAUG TISs (left panel) and others including the AUG TISs in 3’UTRs, non-coding RNAs and de novo-assembled transcripts (right panel). Spearman’s rank correlation coefficient (rho) and the corresponding p-values were shown.

Next, we explored the ability of these models to predict the translation potential for triplets from the sequences of interest. Focusing on *Solyc02g023990*, which contains a novel CDS-AUG TIS driving the translation of a mitochondria-localized protein (**Fig. 3B**) (Li & Liu, 2020), the model identified an AUG triplet within the CDS with a prediction score > 0.93 (orange peak in “WT” plot in **Fig. 3A**). This prediction aligns with experimental evidence showing higher LTM read signals at the same site (“LTM” plot in **Fig. 3A**). When the CDS-AUG TIS was mutated to CUU, the prediction score dropped, corresponding to the abolished protein translation and diminished mitochondrial localization signals (“mt” in **Fig. 3A** and **3B**). For *Solyc03g096920*, the model identified two potential TISs in the 5’UTR with prediction scores > 0.96, one of which matched the experimentally identified 5’UTR-AUG TIS (orange peak in “WT” in **Fig. 3C**) (Wu, TY *et al*., 2024). Mutating the CU-rich sequence upstream of this 5’UTR-AUG TIS led to reduced prediction scores (“mCU” in **Fig. 3C**), consistent with the importance of CU-rich sequence features for TIS prediction (**Fig. 2D**) and reduced protein production (**Fig. 3D**) (Wu, TY *et al*., 2024). Together, these results demonstrated the robustness of the prediction models in identifying TISs across datasets, species, and even at the single gene level. By analyzing sequence features, the *TISCalling* framework provides a powerful sequences-aware tool for identifying potential AUG and non-AUG TISs.

### *TISCalling* framework effectively profiles potential TISs in viral transcripts

Viruses rely on host translational machinery for protein synthesis, intriguing us to investigate whether TIS prediction models developed for hosts could be applied to viral genomes. Indeed, the TIS prediction models derived from Arabidopsis successfully predicted TISs reported in begomoviruses (Chiu *et al*., 2022) (**Fig. 4A**, *At* and *Sl*), despite slightly lower prediction performance compared to that in the host (**Fig. 2**). Similarly, human and mouse TIS models, especially the 5’UTR-nonAUG and CDS-AUG models, predicted TISs reported in human cytomegalovirus and SARS-CoV-2 genomes (Stern-Ginossar *et al*., 2012; Finkel *et al*., 2021)(right two heatmaps in **Fig. 4A; Supporting Dataset S2** for full results). These results highlight the versatility of the *TISCalling* framework in profiling TISs in the viral genomes using models trained on host TISs.

**Figure 4.**
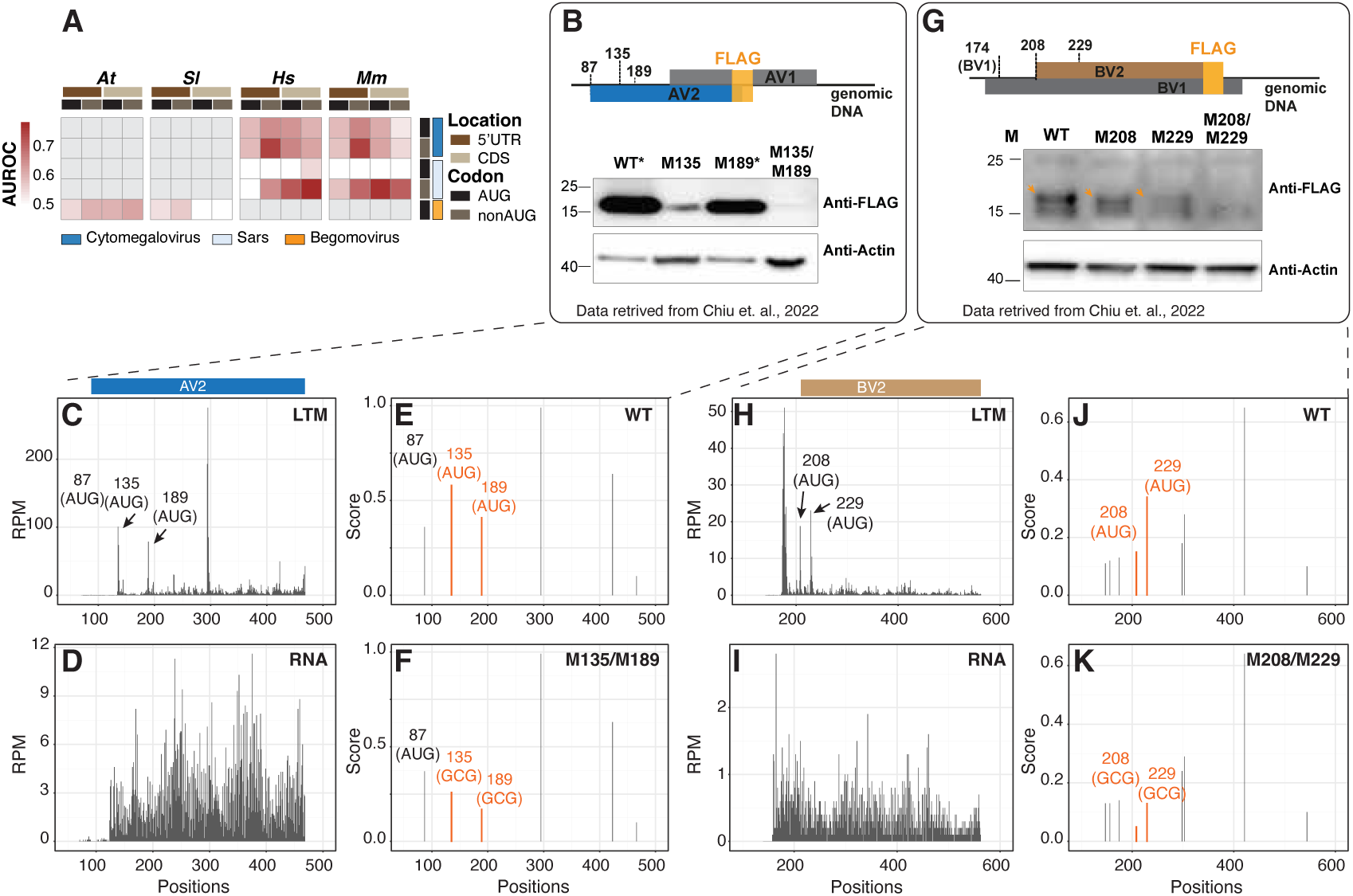
Viral TISs can be predicted using host TIS predictive models. (A) The prediction performance (AUROC scores) of using the four TIS type-generated prediction models from tomato (Sl), Arabidopsis (At), human (Hs), and mouse (Mm) (as indicated in Fig. 2A) to predict the TISs along viral genomes. The viral TISs were derived from different public datasets of plant begomoviruses, human cytomegalovirus and SARS-CoV-2 viruses (Stern-Ginossar et al., 2012; Finkel et al., 2021; Chiu et al., 2022) (See Supporting Dataset S1 for the TIS dataset information). (B) Expression of AV2-FLAG proteins during virus infections with a plant begomovirus infectious clone, containing a FLAG protein insertion at the C-terminus of AV2. Results derived from a previous study (Chiu et al., 2022) were shown for begomovirus infectious clone with WT sequences or specific mutations at nucleotide 135 (M135; M135AUG->GCG) and/or 189 (M189AUG->GCG). In the gene models (top), the black line indicates viral genomic DNAs, boxes represent annotated CDS of viral AV1 and AV2 genes, and the inserted FLAG region. The annotated TIS at position 87 and novel TISs at positions 135 and 189 of the AV2 gene were highlighted. *: WT and M189 samples were diluted 20-fold to equalize the protein abundance for presentation clarity. (C-F) As shown in Fig. 3A, but for the read density at LTM (C) and mRNA (D) levels along viral genomes. The Arabidopsis CDS-AUG TIS model-generated prediction scores are based on WT (E) and M135/M189 mutant (F) sequences. (G) As described in (B), but showing FLAG-tagged BV2 proteins (orange arrows) expressed during virus infection. In the gene model (top), the BV1 TIS at position 174 and two novel AUG TISs of BV2 at positions 208 and 229 were highlighted. Results were shown for infectious clones with the WT sequence or the specific mutations at positions 208 (M208; M208AUG->GCG) and/or 229 (M229; M229AUG->GCG). (H-J) As shown in (C-F), but presenting the results along the BV2-flanking regions of the viral genomes. The results of immunoblotting and LTM plots were derived from a previous study (Chiu et al., 2022).

At the single gene level, previous studies identified novel TIS activities at positions 135 and 189 of *AV2* genes and at positions 208 and 229 of *BV2* genes on begomovirus genomes through LTM-seq analyses (black arrows in **Fig. 4C,H**) (Chiu *et al*., 2022). Consistent with these findings, the *TISCalling* framework assigned relatively high prediction scores to most of these TIS sites (0.58, 0.41, 0.15, 0.34 for each site, respectively) (orange peaks in **Fig. 4E,J**). Mutating these TIS sites (AUG → GCG) in *AV2* and *BV2* genes resulted in lower prediction scores (red peaks in **Fig. 4E,F,J,K, Supporting Dataset S4** for full results), and correlated with reduced protein expression due to the loss of initiation activities at these sites (**Fig. 4B,G**). These findings demonstrate that the *TISCalling* framework, developed for host plants and mammals, can effectively profile potential TISs in viral transcripts, providing insights into translation regulation in viral genomes.

### *TISCalling* framework identifies potential TISs in stress-responsive genes and ncRNAs

The characterization of upstream open reading frames (uORFs), key regulatory elements for the translation of stress-related genes (von Arnim *et al*., 2014; Urquidi Camacho *et al*., 2020; Zhang *et al*., 2020; Wu, TY *et al*., 2024), remains limited due to annotation bias and the lack of translational evidence (Couso, 2015; Hsu & Benfey, 2018). Using the *TISCalling* framework, we identified a predicted AUG TIS in the 5’UTR of Arabidopsis *Heat Shock Transcription Factor B1* (*AtHSFB1*; **Fig. 5C**), corresponding to the TIS of the known uORF1 (Pajerowska-Mukhtar *et al*., 2012; Zhu *et al*., 2012). This TIS/uORF1 did not show significant CHX nor LTM signals in Arabidopsis suspension cells or heat stress (HS)-treated pollen cells (**Fig. 5A,B**) (Willems *et al*., 2017; Poidevin *et al*., 2021), likely due to tissue-specific gene expressions. Further analysis of the TIS-prediction scores revealed three additional potential TISs: one at an AUG codon and two at AUA and CUG codons (orange peaks in **Fig. 5C**). These sites displayed strong LTM and CHX signals but were previously unannotated (**Fig. 5A,B**). Similarly, for other heat-responsive genes, *Heat Shock Transcription Factor B2B* (*AtHSFB2B*) and *Heat Shock protein 70* (*AtHSP70*), the *TISCalling* models predicted multiple AUG and non-AUG TISs (**Fig. 5D-5F; Supplemental Fig. S2**;). Some of these sites exhibited weak LTM/CHX signals in Arabidopsis suspension cells and pollen cells (Willems *et al*., 2017; Poidevin *et al*., 2021), likely due to sequencing depth (**Fig. 5D,E; Supplemental Fig. S2A,B**). The tomato orthologous of these HSF genes also exhibited potential TISs in their 5’UTRs (**Supplemental Fig. S3**), underscoring the role of these uORFs-associated TISs in regulating these heat stress responsive genes. Notably, the non-AUG TIS model identified several potential TIS sites supported by LTM/CHX signals (orange peaks in **Fig. 5C,F**). These findings highlight the utility of *TISCalling* in uncovering previously uncharacterized TISs and uORFs, expanding the horizon of translational regulation in stress responses.

**Figure 5.**
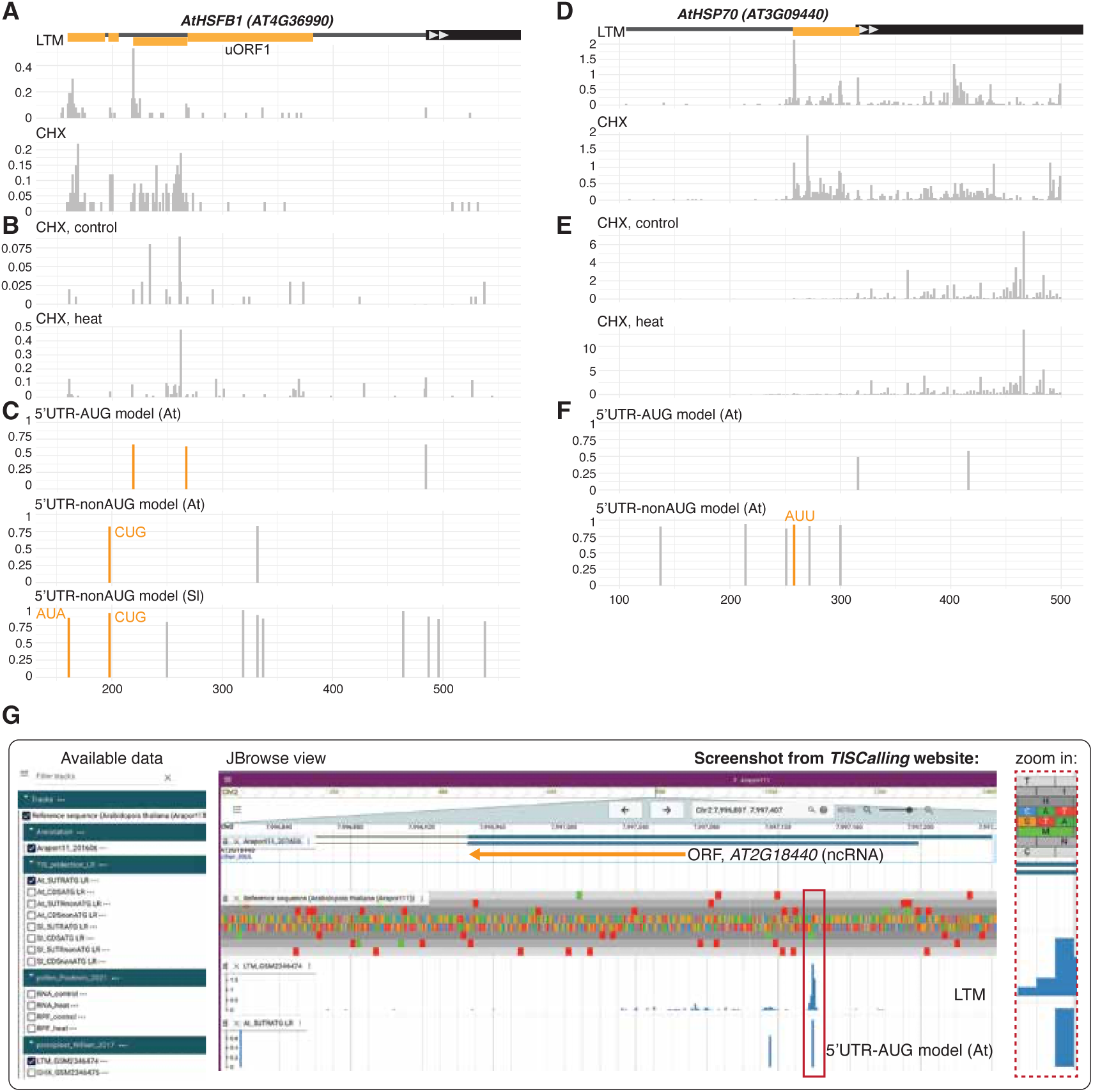
Case study: the potential TISs in plant heat stress-responsive and non-coding RNA genes using TISCalling. (A-C) As shown in Fig. 3A, but for the LTM and CHX signals from Arabidopsis suspension cells (A) (Willems et al., 2017), the CHX-seq signals from heated-treated Arabidopsis pollens (Poidevin et al., 2021) (B) and for the prediction scores generated from Arabidopsis (At) and tomato (Sl) TIS-prediction models (C) along AtHSFB1 transcript. The triplets with high prediction scores and LTM/CHX signals were marked in orange, inferring a potential TIS. In the gene models (top), the black line indicates mRNAs and black and orange boxes represent annotated CDS and the putative ORFs initiation from the indicated potential TISs (orange peaks). (D-F) As indicated in (A-C), but for the potential TISs (orange peaks) and their corresponding ORFs (orange boxes) on AtHSP70. (G) As shown in panels (A-C), this screenshot illustrates the visualization of the predicted AUG-TISs, the LTM signals (red box and the zoon in), and the corresponding ORFs along non-coding RNA (ncRNA) of AT2G18440.1 (orange arrows) using the web-based TISCalling tool.

Non-coding RNAs (ncRNAs), once classified as no coding potential, are now known to encode small peptides and proteins in plants (Sruthi *et al*., 2022; Wang *et al*., 2023). Using the *TISCalling* framework, we identified potential TISs initiating ORFs in ncRNAs, such as *AT2G18440.1*, with high prediction scores (red box in **Fig. 5G**) supported by LTM and CHX signals and previous findings (Hsu *et al*., 2016). The integration of TIS prediction scores with ribosome profiling datasets highlights the ability of *TISCalling* to systematically profile both annotated and uncharacterized AUG and non-AUG TIS/ORFs in ncRNAs. Together, these results demonstrate the versatility of *TISCalling* in accurately predicting TISs across plant heat-stress-responsive genes and ncRNAs. By incorporating LTM/CHX-based ribosome profiling datasets, the framework provides a powerful tool for uncovering hidden translational activities in protein-coding and non-coding RNAs.

### A web-based *TISCalling* tool visualizing potential TISs and ORFs

To facilitate the explorations of TISs at both genome-wide and single-gene scales, we developed a web-based *TISCalling* tool [https://predict.southerngenomics.org/TIScalling]. This user-friendly JBrowse-based platform integrates precomputed TIS prediction scores with publicly available LTM/CHX-based Ribo-seq datasets for Arabidopsis and tomato, covering both protein-coding and non-coding transcripts. To visualize gene models with corresponding TIS prediction scores and LTM/CHX signals, users only need to 1) input a target gene ID, 2) select the species, 3) choose TIS prediction models and Ribo-seq datasets. The tool supports uploading Ribo-seq datasets for comparison with those from additional experimental conditions. This interactive interface allows users to profile and visualize potential TISs and their corresponding ORFs for target genes of interest (**Fig. 5G**; **Supplemental Fig. S3**). To expand the application of *TISCalling* to diverse species, we also present the framework as a Python-based package. This package provides detailed documentation and an easy-to-run command-line interface for generating custom prediction models, identifying key sequence features, and profiling TISs using user-defined datasets and genomes [https://github.com/yenmr/TIScalling].

Together, *TISCalling*, with its web tool and Python package, provides an efficient and robust approach for profiling predicted AUG and non-AUG TISs. The sequence-aware prediction capabilities, independent from Ribo-seq datasets, and flexibility and expansibility of integrating additional omics data make it a versatile tool for exploring coding regions at both gene and genome scales.

## DISCUSSION

The annotation of protein-coding genes often overlooks the translation events involving short coding regions or initiation sites from AUG and non-AUG codons, leaving important aspects of genomic sequences underexplored. To address this, we propose *TISCalling*, a flexible framework designed to systematically investigate potential TISs along transcripts (**Fig. 1**). This framework features sequence-aware machine learning (ML)-based TIS-prediction models capable of robustly predicting AUG and non-AUG TISs independently of Ribo-seq datasets while identifying key sequence features associated with TIS recognition mechanisms in plants and mammals (**Fig. 1,2**). By analyzing mRNA sequence information, the TIS-prediction models generate prediction scores for AUG and non-AUG triplets, enabling the profiling of potential TISs in plant and viral transcripts (**Fig. 3,4**) and the identification of ORFs in protein-coding and non- coding RNAs (**Fig. 5**). To support the broad applicability, *TISCalling* is available as a user-friendly web tool and a command-line Python package, offering users to build custom TIS prediction models, analyze mRNA sequences across genomes, profile and visualize potential TISs at both single-gene and genome-wide scales (**Fig. 1,5**). While intended as an alternative and complementary approach to established TIS-identification methods in plants (Calviello *et al*., 2016; Reuter *et al*., 2016; Zhang, P *et al*., 2017; Machkovech *et al*., 2019; van der Horst *et al*., 2019; Wu, HL *et al*., 2024a), *TISCalling* aims to provide a sequence-aware platform for profiling potential TISs comprehensively. We believe it can offer novel insights into the mechanisms by which ribosomes recognize AUG and non-AUG TISs across species and potentially shed light on previously uncharacterized sequence features in plants and viruses.

Our TIS-prediction model, built on sequence information, computes a prediction score for potential TISs (**Fig. 1,3-5**). Independent of experimental datasets, this approach complements the Ribo-seq-supported TIS identification methods (i.e., those with significant signals and/or phasing patterns from Ribo-seq datasets) and comparative genomics analyses (Calviello *et al*., 2016; Zhang, P *et al*., 2017; van der Horst *et al*., 2019; Wu, HL *et al*., 2024a) and serves as a stand-alone tool for the research community when experimental datasets are unavailable. By analyzing the likelihood of each triplet to function as a TIS, our model enables the generation of the datasets for all possible AUG and non-AUG ORFs across genomes. Despite overall robustness of the model, a known limitation is the potential for false-positive predictions, particularly in non-AUG models (**Fig. 5**). Nevertheless, the TIS-prediction model in *TISCalling* framework is intended to generate comprehensive datasets of potential TISs/ORFs, which can then be refined by integrating Ribo-seq data. This integration improves sensitivity and resolution, enabling a more accurate assessment of translational activity and the biological functions of genomic and genic regions. To prioritize TIS candidates for downstream validation, the LTM/CHX signals from Ribo-seq analyses should be considered, enabling targeted functional analysis of promising TISs.

The performance of TIS-prediction models varies across different regions of transcripts, reflecting the complexity of translational regulation. The 5’UTR-AUG and -nonAUG models generally outperformed the CDS-AUG and -nonAUG ones (**Fig. 2A**). This discrepancy may be attributed to the complexities of multiple translational regulations within CDSs, such as translation initiation/elongation and ribosome stalling (Merret *et al*., 2015; Benitez-Cantos *et al*., 2020; Collart & Weiss, 2020; Iwakawa *et al*., 2021), which may introduce noises in defining true-positive (TP) and true-negative (TN) TISs. Additionally, our model considers only 100-bp upstream of TISs, which is likely sufficient for scanning ribosomes recognizing TISs in 5’UTRs but may be less suitable for TISs within CDS, where ribosomes scan longer regions from the 5’end to the CDSs of the transcripts. Similarly, predicting TISs in viral genomes presents unique challenges. For viral TISs, the highest prediction performance reached an AUROC of ~0.6, suggesting room for improvement. The current viral TIS predictions rely on models built from host data. However, viruses are known to utilize diverse translational strategies, such as internal ribosome entry site (IRES) and secondary structures, to initiate translation from unconventional sites, maximizing protein diversity of their compact genomes (Lozano & Martinez-Salas, 2015; Miras *et al*., 2017; Jaafar & Kieft, 2019). Incorporating these sequence and structural features along with Ribo-seq-supported viral TISs will refine the development of plant viral TIS-prediction models in the future. The *TISCalling* framework represents a sequence-aware tool, independent of Ribo-seq datasets, designed to profile both AUG and non-AUG initiation sites at the genome and single-gene levels. By integrating prediction models with visualization capabilities, *TISCalling* offers a versatile approach for exploring translational initiation mechanisms across diverse genomic regions. The framework is accessible through a command-line Python package on Github and a user-friendly JBrowse tool to visualize and profile potential TISs across plants and viruses with simplicity, flexibility, and expandability. This advancement opens new opportunities for identifying potential AUG- and non-AUG-initiated ORFs across genomes. Moreover, it enhances our ability to uncover the biological significance of previously unexplored genomic regions and offers insights into the mechanisms driving translation initiation in plant and viral species. With its flexibility to integrate additional omics datasets and expand to other species, *TISCalling* serves as a valuable resource for advancing genome annotation and understanding translational regulation in diverse contexts.

## ACKNOWLEDGMENTS

We thank the core facilities of Transgenic Plant Laboratory and AS-BCST Bioinformatics Core for high-performance computing services. We acknowledge Biorender (https://www.biorender.com) for creating **Fig. 1A**. This research was financially supported by grants of AS-CDA-111-L06 to Ming-Jung Liu and AS-CDA-113-L03 to Ting-Ying Wu.

## COMPETING INTEREST STATEMENT

The authors declare that they have no conflict of interest.

## AUTHOR CONTRIBUTION

M.-R. Y. performed the computational analyses and developed the command-line-based and web-based tools. C.-Y. C. developed the command-line-based and web-based tools and revised the paper. T.-Y. W. and M.-J. L. conceived/designed the research, interpreted the results, and wrote the paper.

## DATA AVAILABILITY

The LTM- and CHX-based Ribo-seq in Arabidopsis (suspension cells), Arabidopsis pollens and tomato were retrieved from NCBI Gene Expression Omnibus (GEO; https://www.ncbi.nlm.nih.gov/geo/) under accession number GSE145795, GSE88790, GSE143311

## SUPPLEMENTAL MATERIAL

**Supplemental Figure 1. The workflow for generating sequences features and prediction models in *TISCalling***

Ten sets of the balanced True positive (TP)/True Negative(TN) translation initiation site (TIS) datasets were generated via random sampling and used for feature selections and model generation, separately. The sequence features focused on the 200-bp regions centered on target TISs and were significantly different between TP and TN TIS datasets (Mann-Whitney test, *p*< 0.01) while showing low correlation with other features (Pearson correlation, *r* < 0.7). Support Vector Machine (SVM) and Logistic Regression (LR) algorithms were applied to generate the TIS prediction models using the selected sequence feature sets to distinguish between TP and TN TISs. This pipeline was adapted from a previous study with minor modifications (Reuter *et al*., 2016) (see Methods for more details). The matrix of F1 scores, Matthews Correlation Coefficient (MCC), and Area Under the Receiver Operating Characteristic Curve (AUROC) were for evaluating the model performance.

**Supplemental Figure 2. Case study: identifying potential TISs in Arabidopsis heat stress-responsive genes using *TISCalling*.**

(A-C) The LTM and CHX plots showed the read density (reads per million mapped reads; RPM) of LTM- and CHX-treated Ribo-seq from Arabidopsis suspension cells (Willems *et al*., 2017) (A) and from heated-treated Arabidopsis pollens (Poidevin *et al*., 2021) (B) across *AtHSFB2B (AT4G11660)* transcript. The prediction scores of a given triplet along the transcript were generated via Arabidopsis AUG- and non-AUG TIS models and were based on the sequences of the triplet and its flanking 200-bp regions (C). In the gene model (top), light and dark gray boxes indicate UTRs and annotated CDSs, while orange boxes indicate the putative upstream ORF initiated from the prediction TISs (orange peaks in (C)).

**Supplemental Figure 3. Case study: identifying potential TISs in tomato heat stress-responsive genes using *TISCalling*.**

(A-C) As indicated in **Supplemental Fig. S2A-C**, but for a screenshot illustrating the visualization of the predicted UUG-, AUU, and CUG-TISs and the LTM signals (red boxes and the zoom in) and the corresponding ORFs (orange arrows) across the tomato orthologs of three heat-stress responsive genes via the web-based *TISCalling* tool. The LTM signals were from tomato leaves (Li & Liu, 2020).

**Supporting Dataset 1. The translation initiation site (TIS) datasets used in this study.**

**Supporting Dataset 2. Model performance shown in terms of F1 scores, Matthews Correlation Coefficient (MCC), and Area Under the Receiver Operating Characteristic Curve (AUROC).**

**Supporting Dataset 3. The feature information of TIS-prediction models.**

**Supporting Dataset 4. The TIS prediction scores along the genome of Tomato yellow leaf curl Thailand virus (TYLCTHV, genus begomovirus).**

## Notes

### Competing Interest Statement

The authors have declared no competing interest.

