## Supporting_figures for "*TISCalling:* Leveraging Machine Learning to Identify Translational Initiation Sites in Plants and Viruses"

#### Supplemental Figure S1

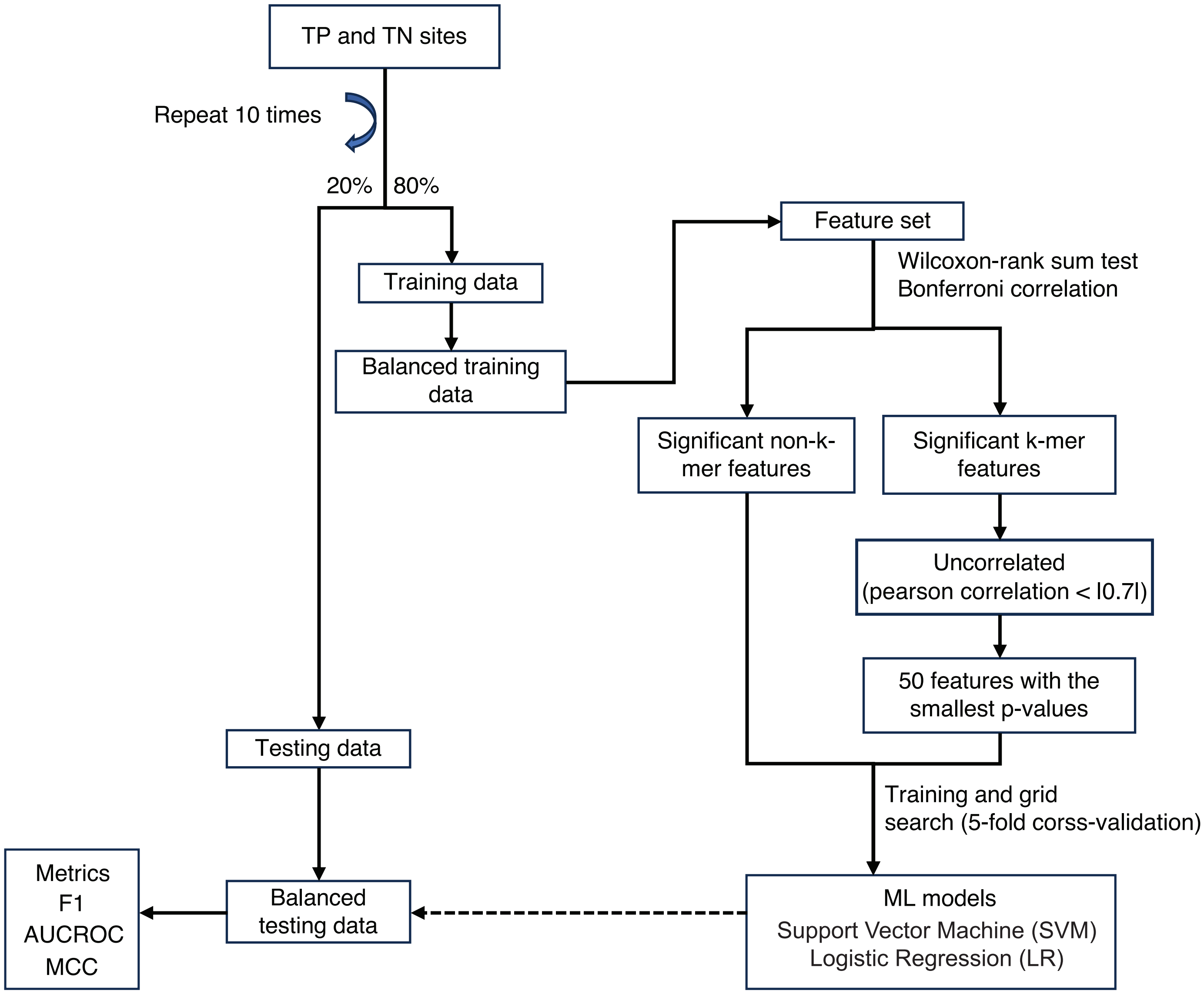


**Supplemental Figure 1. The workflow for generating sequences features and prediction models in TISCalling**

Ten sets of the balanced True positive (TP)/True Negative(TN) translation initiation site (TIS) datasets were generated via random sampling and used for feature selections and model generation, separately. The sequence features focused on the 200-bp regions centered on target TISs and were significantly different between TP and TN TIS datasets (Mann-Whitney test, $p < 0.01$) while showing low correlation with other features (Pearson correlation, $r < 0.7$). Support Vector Machine (SVM) and Logistic Regression (LR) algorithms were applied to generate the TIS prediction models using the selected sequence feature sets to distinguish between TP and TN TISs. This pipeline was adapted from a previous study with minor modifications (Reuter et al., 2016) (see Methods for more details). The matrix of F1 scores, Matthews Correlation Coefficient (MCC), and Area Under the Receiver Operating Characteristic Curve (AUROC) were for evaluating the model performance.

### Supplemental Figure S2

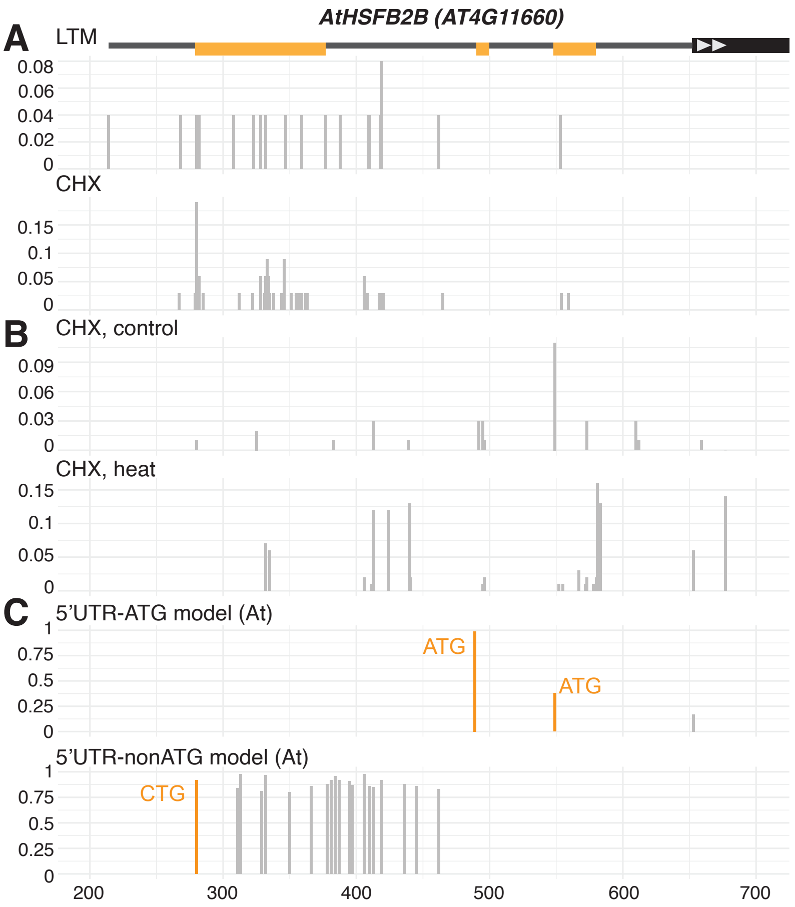


**Supplemental Figure 2. Case study: identifying potential TISs in Arabidopsis heat stress-responsive genes using TISCalling.**
(A-C) The LTM and CHX plots showed the read density (reads per million mapped reads; RPM) of LTM- and CHX-treated Ribo-seq from Arabidopsis suspension cells (Willems et al., 2017) (A) and from heated-treated Arabidopsis pollens (Poidevin et al., 2021) (B) across AtHSFB2B (AT4G11660) transcript. The prediction scores of a given triplet along the transcript were generated via Arabidopsis AUG- and non-AUG TIS models and were based on the sequences of the triplet and its flanking 200-bp regions (C). In the gene model (top), light and dark gray boxes indicate UTRs and annotated CDSs, while orange boxes indicate the putative upstream ORF initiated from the prediction TISs (orange peaks in (C)).

### Supplemental Figure S3

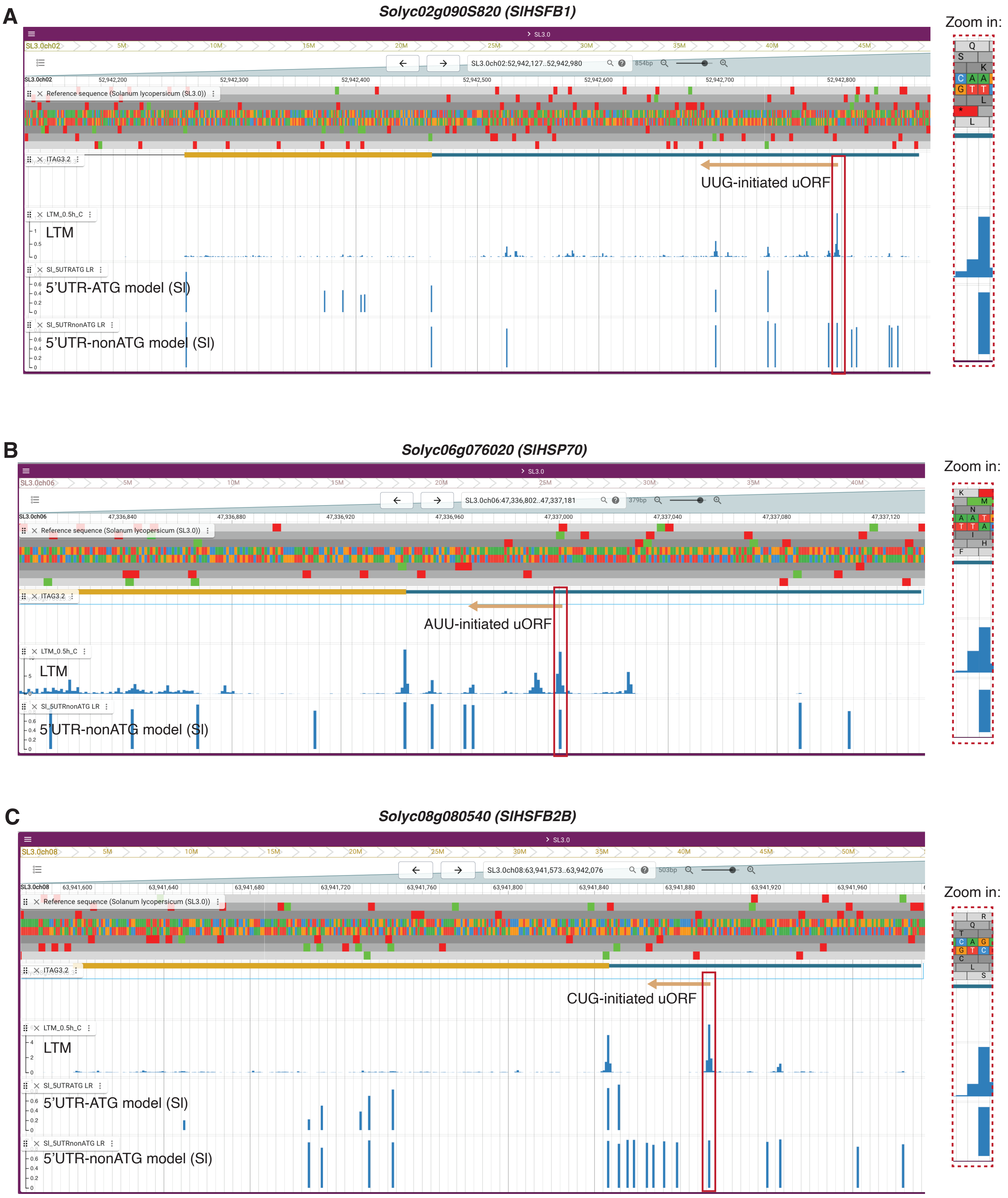


**Supplemental Figure 3. Case study: identifying potential TISs in tomato heat stress-responsive genes using TISCalling.**
(A-C) As indicated in Supplemental Fig. S2A-C, but for a screenshot illustrating the visualization of the predicted UUG-, AUU, and CUG-TISs and the LTM signals (red boxes and the zoom in) and the corresponding ORFs (orange arrows) across the tomato orthologs of three heat-stress responsive genes via the web-based TISCalling tool. The LTM signals were from tomato leaves (Li & Liu, 2020).
